# The systemic JIA synovial fluid environment supports development and prevalence of specific inflammatory T helper cell phenotypes

**DOI:** 10.1101/2025.07.01.661738

**Authors:** Susanne Schleifenbaum, Aljoscha Swoboda, Johannes Dirks, Claudia Bracaglia, Ivan Caiello, Tanja Hinze, Sven Hardt, Giusi Prencipe, Giusyda Tarantino, Manuela Pardeo, Sabrina Fuehner, Carolin Park, Claas Hinze, Helmut Wittkowski, Daniel Windschall, Dirk Foell, Henner Morbach, Christoph Kessel

**Author notes:** **Corresponding author:** Dr. Christoph Kessel, Department of Pediatric Rheumatology and Immunology, University Children’s Hospital Muenster, Domagkstr. 3, 48149 Muenster, Germany.

## Abstract

**Objective:** The potential involvement of adaptive immunity in systemic juvenile idiopathic arthritis (sJIA) pathophysiology remains an intriguing question. Here, we investigated whether and how the inflammatory environment in sJIA versus JIA synovial fluid (SF) may differentially impact T helper (Th) cell polarization and activation.

**Methods:** SF samples from sJIA and JIA patients (both n=7) were tested in various cell culture setups, with or without recombinant cytokines or cytokine-blocking drugs, to assess their effects on healthy donor Th cell activation. We analyzed cellular surface marker, transcription factor, and effector molecule expression using flow cytometry, Luminex, ELISA, and qRT-PCR.

**Results:** Both sJIA and JIA SF revealed highly pro-inflammatory profiles. Compared to JIA, sJIA SF demonstrated markedly elevated IL-1β, IL-18, GM-CSF, S100A9, and MPO levels, while JIA SF showed trends toward higher soluble FasL and IL-17A concentrations. Notably, sJIA SF significantly increased CD4 T cell ICOS expression and expanded CXCR3^pos^CCR6^pos^ Th cells, whereas JIA SF favored expansion of CXCR3^neg^CCR6^pos^ Th cells and CCR6^pos^ Th cell expansion was sensitive to IL-1 blockade. Systemic JIA SF selectively sustained a IFNγ/IL-21 expressing T peripheral helper (Tph) phenotype, particularly associated with IL-1β, IL-18, and GM-CSF SF levels. Spiking JIA SF with a cocktail of these cytokines recapitulated some T cellular phenotypic features observed in sJIA SF cultures.

**Conclusion:** JIA and sJIA SF drive distinct Th cell polarization, including differential and sustained Tf/ph cell activation. These findings complement our earlier observations in sJIA peripheral blood and demonstrate the impact of the SF inflammatory matrix on immune cell activation.

**What is already known on this topic:**

- Systemic juvenile idiopathic arthritis (sJIA, Still’s Disease) is initially hallmarked by innate immunity driving systemic inflammation
- With further disease course sJIA can progress to chronic destructive arthritis with several studies suggesting an involvement of adaptive immunity in this process.
- In a previous study we linked T follicular/peripheral helper (Tfh/Tph) cells in sJIA patients’ blood to arthritis and self-reactive antibody signatures in patients with longer disease duration

**What this study adds:**

- Our study demonstrates that sJIA versus JIA synovial fluid (SF) can drive differential T helper cell (Th cell) polarization and activation.
- Our data imply that sJIA SF promotes the self-sustenance of Th cells with a Tph phenotype characterized by IFNγ, IL-21 and c-MAF expression.
- IL-1β, IL-18, and GM-CSF levels in sJIA SF can be associated with Tph perseverance, and recapitulate sJIA Tph features when spiked into JIA SF.

**How this study might affect research, practice or policy:**

- The present data complement our earlier observations from sJIA peripheral blood and strengthen the biphasic model hypothesis regarding sJIA progression
- While both JIA and sJIA patients can develop clinically similar arthritis, our data demonstrate how the respective local inflammatory environments can differentially impact and shape T cell immunity.
- Our data suggest a combined and early targeting of both IL-1β and IL-18 in sJIA may be effective in preventing the generation of inflammatory T cell subsets with the potential to drive chronic arthritis through a joint-localized, adaptive immune response.

## Introduction

Juvenile idiopathic arthritis (JIA) is the most common chronic rheumatic disease among children and is defined as arthritis of unknown origin with onset before the age of 16 years. Clinical presentation can involve arthralgia, synovitis, arthritis, psoriasis or enthesitis and – next to clinical laboratory parameter – defines disease subtypes according to the current International League of Associations for Rheumatology (ILAR) classification (1, 2).

Oligoarticular and polyarticular JIA represent around 70-80% of cases. Oligoarticular JIA affects fewer than five joints, while polyarticular JIA involves five or more joints within the first six months following onset. These JIA subtypes are thought to be primarily driven by adaptive immunity, characterized by T and B cell activation, autoantibody production (ANA, RF, ACPA), and abnormal B cell receptor rearrangements, which suggests a significant autoimmune disease component in a majority of patients.(2, 3)

In contrast, systemic JIA (sJIA) accounts for 10-20% of JIA cases and exhibits distinct features such as spiking fevers, rash, hepatosplenomegaly, lymphadenopathy, and serositis(4). Today, sJIA together with Adult Onset Still’s Disease (AOSD) are considered as one inflammatory continuum and are labeled as Still’s Disease (SD)(5), yet most data on specific molecular pathomechanisms have been acquired in context with sJIA. In sJIA arthritis may be minimal or absent at onset, but disease can progress to chronic destructive joint inflammation if untreated or treatment-refractory(6, 7). SD is primarily associated with excessive innate immune activation, resulting in disease features also prevalent in IL-1-driven autoinflammatory fever syndromes. This is largely supported by significant improvements in disease outcomes upon therapeutic IL-1 blockade(8), particularly in the early systemic disease phase (9, 10).

A proposed biphasic model of SD pathogenesis suggests that dysregulated innate immunity in early disease may prime downstream adaptive responses, which can drive chronic inflammation and destructive arthritis in the further disease course(7, 11). Genetic associations with HLA-DRB1*11(12) together with evidence for IL-1-driven Th17 differentiation(13) and altered phenotypes and frequencies of γδT, and natural killer (NK) cells(14, 15) as immune cells bridging innate and adaptive immunity may indeed suggest an autoimmune component to sJIA(7, 11, 16).

In earlier work, we found that naïve peripheral Th cells in sJIA patients preferentially differentiated into T peripheral/follicular helper (Tph/Tfh) cells(17). Both Tph and Tfh subsets are essential for supporting B cellular antibody production in secondary lymphoid organs (Tfh cells) or within inflamed tissues (Tph cells). Thus, particularly Tph cells have been linked to the pathology of autoimmune conditions in rheumatoid arthritis(18) or systemic lupus erythematosus(19) but also JIA(20, 21).

While our prior analysis was limited to peripheral blood samples, in the present study, we had the unique opportunity to investigate the inflammatory conditioning in the sJIA synovium as a major site of tissue born inflammation. In contrast to JIA patients, sJIA patients are only rarely subjected to intra-articular treatment procedures, which can give access to synovial fluid (SF). Our findings reveal that, compared to JIA SF, the sJIA SF inflammatory matrix triggers and maintains specific Th cell polarization patterns, including sustaining Tph phenotypes. We further linked some of these observations to the specific SF inflammatory mediator profiles and, by spiking specific recombinant cytokines into JIA SF, we were able to partially recapitulate some of the Tph-associated phenotypic features observed in sJIA .Altogether, our data significantly extend our previous observations from the sJIA periphery(17) and highlight the power of the SF inflammatory matrix in shaping T cell immunity.

## Methods

SF of sJIA as well as oligo-and polyarticular JIA patients (hereafter referred to as JIA; n=7 each, **table 1**) was obtained during routine joint puncture and aspiration and used in different cell stimulation experiments. All respective experimental procedures and data analysis are detailed in the supplementary methods section. Synovial cells from the identical sJIA patients as reported herein, but an independent cohort of JIA patients were analyzed in a paralleling study(22).

**Table 1.** Study population of SF donors

|  | **sJIA**<br>(n=7) | **JIA**<br>(n=7) |
| --- | --- | --- |
| **Female**, number (%) | 4 (57) | 6 (86) |
| **Age at onset, (month)**, median (range) | 54 (19-180) | 60 (24-144) |
| **Age at sampling, (month)**, median (range) | 128 (29-300) | 96 (60-192) |
| **Disease duration (month)**, median (range) | 44.3 (9.7-120) | 36 (1-156) |
| **oligoarticular JIA**, number (%) | --- | 4 (57.1) |
| **polyarticular JIA**, number (%) | --- | 3 (42.8) |
| **Treatment** |  |  |
| **Treatment naïve**, number (%) | 0 (0.0) | 3 (42.8) |
| **Glucocorticoids**, number (%) | 2 (28.6) | 0 (0.0) |
| **MTX**, number (%) | 2 (28.6) | 3 (42.9) |
| **TNF-α inhibition**, number (%) | 0 (0.0) | 1 (14.3) |
| **IL-6 inhibition**, number (%) | 5 (71.4) | 1 (14.3) |
| **Clinical laboratory data** |  |  |
| **CRP** (mg/dL), median (range) | 1.6 (<0.05-14.5) | 0.14 (<0.05-0.39) |
| **ESR** (mm/h), median (range) | 5 (2-51) | 5 (2-16) |
| **ANA ≥ 1:160**, number (%) | 1 (12.5) | 7 (100%) |

## Results

### JIA and sJIA SF triggers differential Th cell activation and subset expansion

Initially, we aimed to understand the impact of a sJIA versus JIA SF inflammatory matrix on T cell activation. Therefore, peripheral blood mononuclear cells (PBMCs) freshly isolated from a single healthy donor were cultured with T cell receptor activators (anti-CD3/CD28) in the presence or absence of medium conditioned with (3% or 6%) JIA or sJIA SF. In addition, drugs to inhibit IL-6R signaling (tocilizumab) or IL-1R signaling (recombinant IL-1Ra: anakinra; anti-IL-1β antibody: canakinumab) were added to cultures, or those were left untreated. After five days of culture, cells were analyzed by flow cytometry (**Fig. 1A, B**).

**Figure 1.**
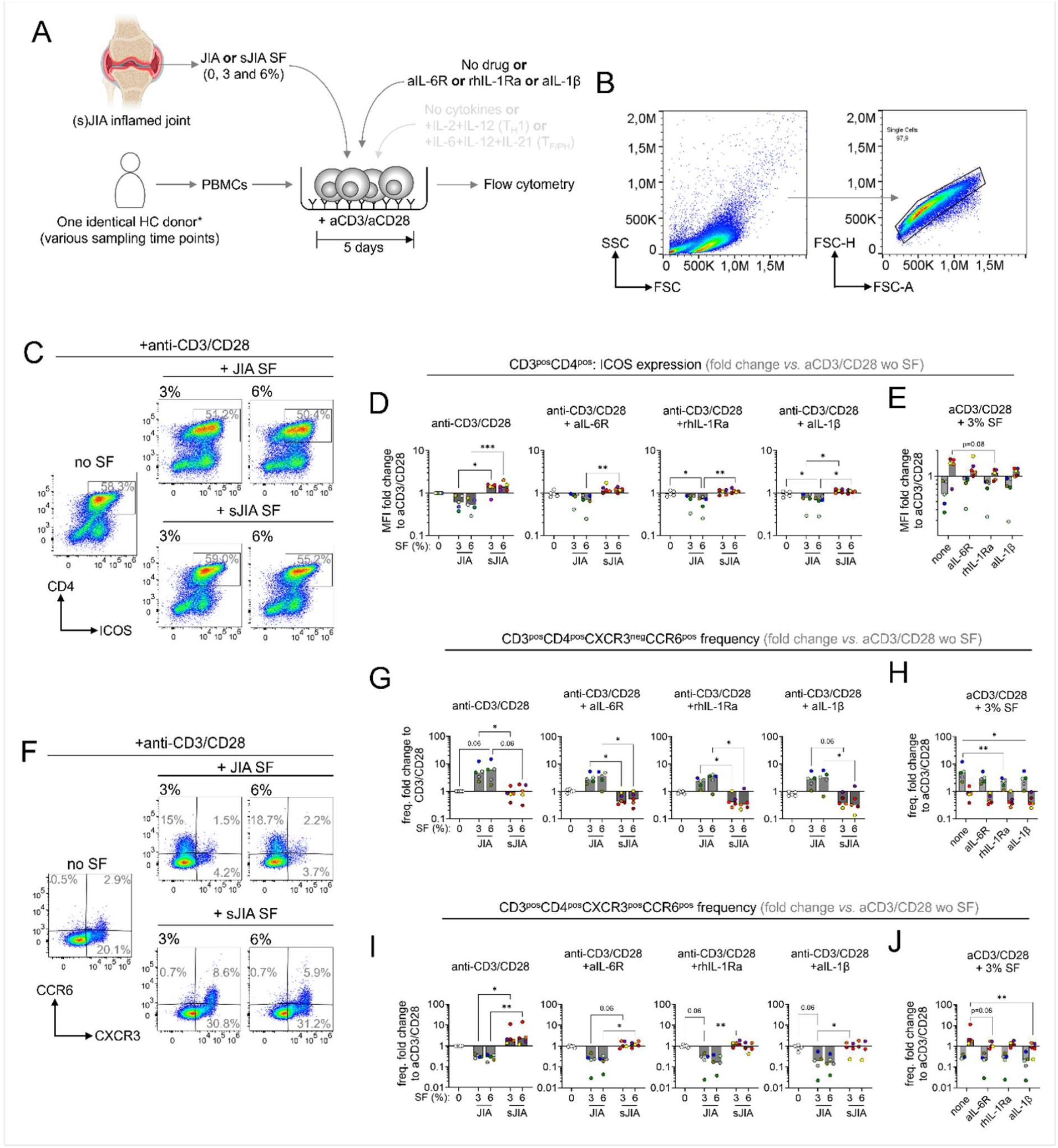
Differential Th cell activation and subset expansion upon HD PBMC culture in JIA or sJIA SF conditioned medium. (**A**) Experimental setup. (**B**) Exemplary gating strategy of single cells following five days of culture. (**C**) Exemplary pseudocolor flow cytometry plots of CD4 T cellular ICOS expression following five days of HD PBMC culture with anti-CD3/CD28 stimulation and JIA or sJIA SF (both 3 and 6%) conditioned medium. (**D**, **E**) Cumulative data (n=5 independent JIA SF donors; n=5 sJIA independent SF donors) of ICOS expression fold change induced by cell cultures as described in **C**), with or without additional *in vitro* drug treatment (anti-IL-6R, tocilizumab; recombinant human IL-1Ra, anakinra; anti-IL-1b, canakinumab). (**D**, **E**) Changes in expression were calculated as fold change compared to mean fluorescence intensities of cells stimulated with only anti-CD3/CD28. (**F**) Exemplary pseudocolor flow cytometry plots of CCR6 and CXCR3 expressing CD4 T cells following five days of HD PBMC culture with anti-CD3/CD28 stimulation and JIA or sJIA SF (both 3 and 6%) conditioned medium. (**G**-**J**) Cumulative data (n=5 independent JIA SF donors; n=5 sJIA independent SF donors) of CXCR3^neg^CCR6^pos^ (**G**, **H**) and CXCR3^pos^CCR6^pos^ (**I**, **J**) fold change induced by cell cultures as described above, with or without indicated *in vitro* drug treatment. (**G**-**J**) Changes in population frequencies were calculated based on fold change of absolute cell numbers in total single cells, compared to cells stimulated with anti-CD3/CD28 alone. All data were analyzed by Friedman test for multiple paired non-parametric observations, followed by Dunn’s multiple comparison test. * = p<0.05, ** = p<0.01; *** = p<0.001

In these experiments we initially observed no difference in Th cell expansion when cells were cultured in either JIA or sJIA SF conditioned medium (**Fig. S1A, B**). Yet, we noted a significant upregulation of CD4 T cellular ICOS expression when cells were cultured in sJIA SF conditioned medium. Conversely, JIA SF induced a concentration dependent decrease in ICOS expression levels (**Fig 1C, D** left panel). Throughout, drug treatment resulted in a partial reduction of these effects, though not reaching statistical significance (**Fig. 1D, E**; 3% sJIA SF cultures untreated vs anakinra: p=0.08). Interestingly, while JIA SF conditioned medium reduced ICOS expression, it also led to a marked, but non-significant upregulation, of PD-1 expression, compared to cells cultured in sJIA SF (**Fig.S1C, D**). *In vitro* drug treatment did not affect this expression phenotype (**Fig. S1E**).

Next to an increase in PD-1 expression, T cell activation in the context of JIA SF induced a marked expansion of a CXCR3^neg^CCR6^pos^ Th cell subset, which we did not observe in cell cultures with sJIA SF (**Fig. 1F, G**). The expansion in CXCR3^neg^CCR6^pos^ Th cells was significantly reduced by inhibition of IL-1β (canakinumab) and IL-1R1 (anakinra) signaling (**Fig. 1H**). Conversely, sJIA SF promoted a moderate expansion of a CXCR3^pos^CCR6^pos^ Th cell subset, which was rather depleted in JIA SF cultures (**Fig. 1F, I**). Expansion in CXCR3^pos^CCR6^pos^ Th cells was significantly reduced upon *in vitro* treatment with anakinra (**Fig. 1J**). The absence of CXCR3 on CCR6 expressing Th cells subsets has been reported to indicate a Th17 effector memory subset and has thus been predominantly linked with IL-17A expression(23). In our data, T cell activation in the context of JIA SF to induce an expansion of CXCR3^neg^CCR6^pos^IL-17A^pos^ Th cells, which was not observed sJIA SF cell cultures and which was significantly reduced by anakinra treatment in particular (**Fig. S1F**, **G**). No expansion of CXCR3^pos^CCR6^pos^IL-17A^pos^ Th cells, reported to resemble Th1-like Th17 cells(23), was observed in cultures with either JIA or sJIA SF (data not shown).

### Systemic JIA but not JIA SF preserves a recombinant cytokine induced T peripheral helper cell phenotype

Following up on observations of differential Th cellular ICOS and PD-1 expression among cells cultured in sJIA *versus* JIA SF, we aimed to analyze the potential impact of the different inflammatory matrices on Tfh or Tph subsets. To assess this, we cultured PBMCs freshly isolated from a single healthy donor under the same conditions as described above. In addition and line with previous experiments(17), we also spiked cultures with recombinant cytokines (Th1: IL-2+IL-12; Tfh/ph: IL-6+IL-12+IL-21), which we previously observed to drive preferential differentiation of sJIA naïve peripheral Th cells towards a Tf/ph phenotype(17) (**Fig. 2A**).

**Figure 2.**
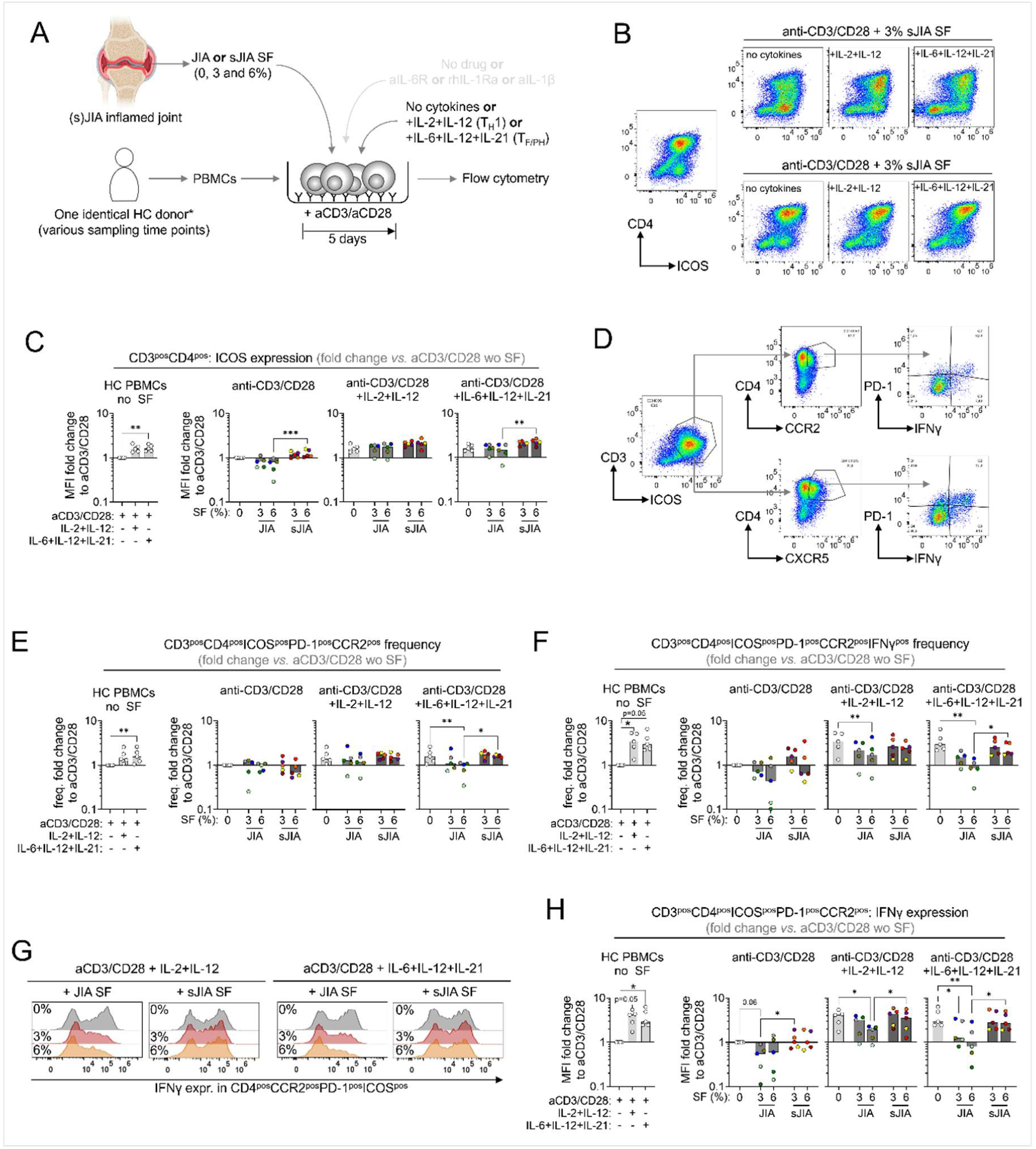
Systemic JIA but not JIA SF preserves a recombinant cytokine induced T peripheral helper cell phenotype. (**A**) Experimental setup. (**B**) Exemplary pseudocolor flow cytometry plots of CD4 T cellular ICOS expression following five days of HD PBMC culture with anti-CD3/CD28 stimulation and JIA or sJIA SF (both 3 and 6%) conditioned medium and addition of indicated recombinant cytokines. (**C**) Cumulative data (n=5 independent JIA SF donors; n=5 sJIA independent SF donors) of ICOS expression fold change induced by cell cultures as illustrated in (**A**), with or without additional recombinant cytokine treatment. (**D**) Exemplary pseudocolor flow cytometry plots and gating strategy of Tph (upper panels: CD3^pos^CD4^pos^CCR2^pos^ICOS^pos^PD-1^pos^) and Tfh (lower panels: CD3^pos^CD4^pos^CXCR5^pos^ICOS^pos^PD-1^pos^) IFNγ expression. (**E**, **F**) Cumulative data (n=5 independent JIA SF donors; n=5 sJIA independent SF donors) of Tph (**E**) and IFNγ expressing Tph expansion (**F**) fold change induced by the indicated cell culture conditions. (**G**, **H**) Exemplary histogram plots (**G**) and cumulative data (n=5 independent JIA SF donors; n=5 sJIA independent SF donors) of Tph IFNγ expression fold change (**H**) induced by the indicated cell culture conditions. (**C**, **H**) Changes in expression were calculated as fold change compared to mean fluorescence intensities of cells stimulated with only anti-CD3/CD28. (**E**, **F**) Changes in population frequencies were calculated based on fold change of absolute cell numbers in total single cells, compared to cells stimulated with anti-CD3/CD28 alone. All data were analyzed by Friedman test for multiple paired non-parametric observations, followed by Dunn’s multiple comparison test. * = p<0.05, ** = p<0.01; *** = p<0.001

Consistent with our results illustrated in **Figure 1** (**C**, **D**), cultures in sJIA SF conditioned medium led to a significant increase of ICOS expression on Th cells compared to cells cultured with JIA SF (**Fig. 2B, C**). The addition of recombinant cytokines alone already increased ICOS expression on cells cultured without SF (**Fig. 2C** left panel), and further enhanced expression on cells cultured in sJIA SF conditioned medium (**Fig. 2C**). Together with JIA SF, recombinant cytokines also increased ICOS expression, but this increase was both less consistent and prominent as observed in the context of sJIA SF, particularly when testing SF in combination with Tfh/Tph polarizing conditions (**Fig. 2C**).

When specifically aiming at analysis of SF inflammatory matrix impact on Tfh (**Fig. 2D** and **Fig. S2**) and Tph subsets (**Fig. 2D-H**), we observed that the addition of Th1, and particularly Tfh polarizing recombinant cytokines promoted a significant expansion of cells with a Tph phenotype (**Fig. 2E**, left panel). This expansion was not affected by the addition of 3% or 6% of sJIA SF to cell cultures (**Fig. 2E**). In contrast, the combination of JIA SF with recombinant cytokine stimulation did rather result in a significant depletion of this cell subset (**Fig. 2E**). Similarly, we also observed a significant increase of ICOS^pos^PD-1^pos^CXCR5^pos^ cells when treating cell cultures with Tfh/Tph polarizing conditions (+ rhIL-6 + rhIL-12 + rhIL-21; **Fig. S2A**, left panel), likely resembling a Tfh phenotype. Beyond this, no significant differences in frequencies of cells with a Tfh phenotype upon addition of either sJIA or JIA SF were observed (**Fig. S2A**).

An even further pronounced negative effect of the JIA SF environment on the recombinant cytokine-induced Tph subset in our cell cultures could be observed when analyzing IFNγ expression, a key effector cytokine of Tph cells as well as a Tfh subset(19, 24, 25). Here we observed that both the addition of Th1 and Tfh polarizing recombinant cytokines resulted in marked expansion of IFNγ-expressing Tph (**Fig. 2F**, left panel) and Tfh cells (**Fig. S2B**). Further, recombinant cytokine treatment in part significantly increased cellular IFNγ expression by Tph (**Fig. 2H**, left panel) and Tfh cells (**Fig. S2D**). Both the frequency of IFNγ-expressing Tph and Tfh cells as well as their IFNγ expression was not affected by the addition of 3% or 6% sJIA SF to cultures (**Fig. 2F-H** and **Fig. S2B-D**). In contrast, when combining JIA SF with particularly Tf/ph polarizing cytokines, this significantly reduced frequencies as well as cytokine expression of IFNγ-producing Tph and Tfh cells (**Fig. 2F-H** and **Fig. S2B-D**).

A reduction of IFNγ-expression upon culture in JIA SF, but not sJIA SF conditioned medium was also observed at secretory and transcriptional levels, as in some of our experiments we also quantified gene expression and cytokine release into culture supernatants (**Figure S3A**, **E**). With the addition of recombinant cytokines, and in line with our flow cytometry data, we consistently observed a depletion in IFNγ-expression, particularly at the transcriptional level (**Figure S3E**). In this analysis we also assessed the expression of IL-21 and CXCL-13 as two Tf/ph hallmark cytokines(25). In experiments stimulating T cells in the presence of sJIA SF, IL-21 expression was increased over levels observed in cell cultures with JIA SF both at gene and protein expression level (**Figure S3B**, **F**). In contrast, *CXCL13, BCL6 and PRDM1* expression was markedly elevated upon T stimulation in cell cultures including JIA SF (**Figure S3G**, **H**, **J**). MAF expression, however, appeared more robust among T cells receiving anti-CD3/CD28 stimulation with or without additional recombinant cytokines in the presence of sJIA SF (**Figure S3I**). Finally, we also assessed IL-17A release into culture supernatants, which was markedly promoted by sJIA SF in all tested cell culture conditions, unlike JIA SF (**Figure S3D**).

### Perseverance of Tf/ph activation by systemic JIA SF associates with synovial inflammatory cytokine levels

The observed differences in activation and propagation of specific inflammatory Th cell subsets depending on an exposure to JIA *versus* sJIA SF environment prompted us to ask, whether we might be able to determine factors directly contributing to the observed Th cell phenotypes. Therefore, we performed a multiplexed analysis of inflammatory mediator expression in JIA and sJIA SF (**Fig. 3A**). While most of the quantified proteins were markedly elevated over healthy control serum levels in both JIA and sJIA SF (**Fig. 3B-D**), several inflammatory mediators (IL-1β, IL-18, GM-CSF, MPO, S100A9 and E-selectin) where significantly overexpressed in sJIA compared to JIA SF (**Fig. 3B**). Conversely, we observed sFasL, IL-17A and sVCAM-1 to be elevated by trend in JIA over sJIA SF (**Fig. 3C**). Beyond, several other inflammatory mediators were elevated in sJIA over JIA SF by trend (**Fig. 3D**).

**Figure 3.**
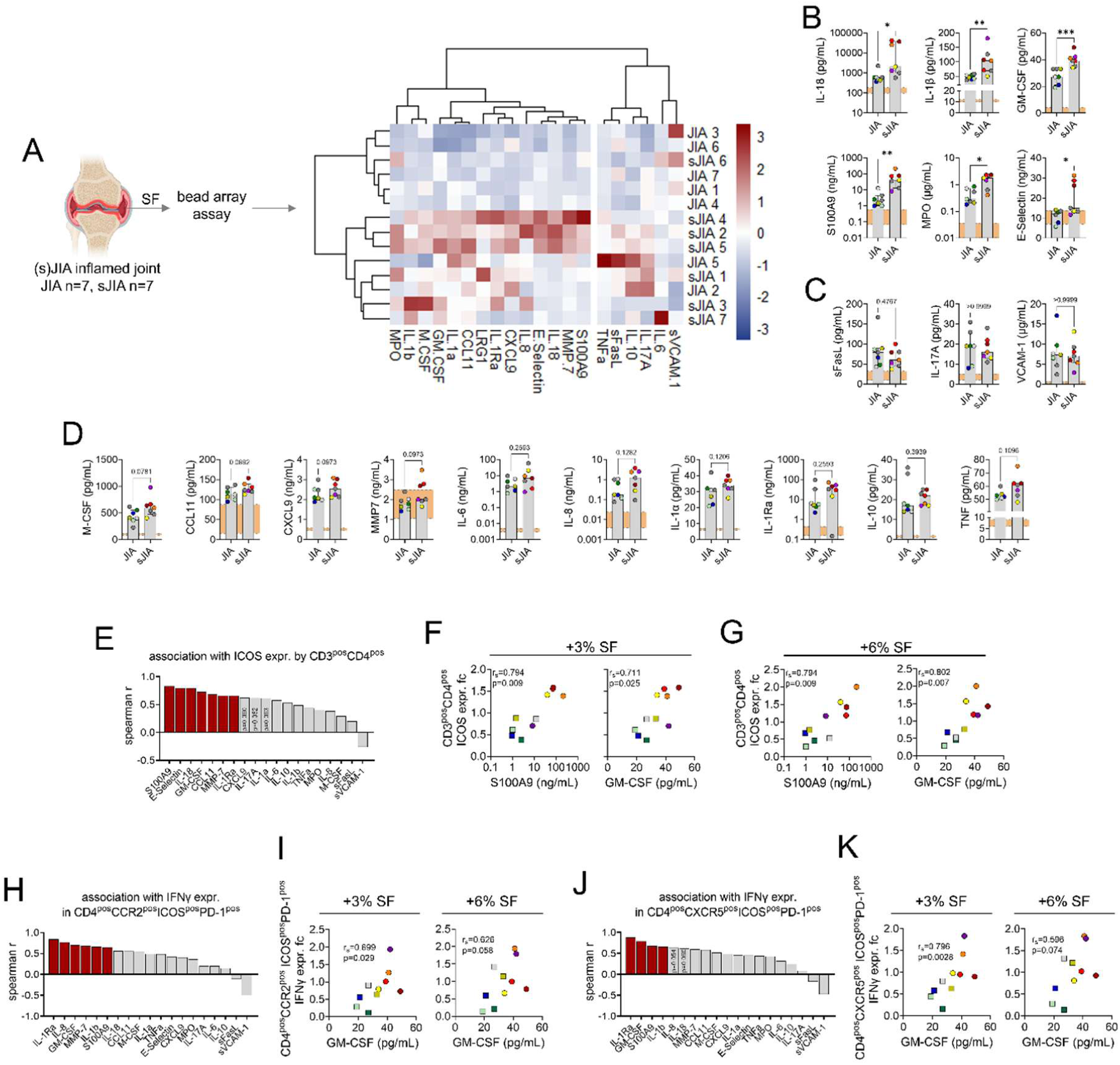
Impact of SF on Tf/ph phenotype maintenance associates with synovial inflammatory cytokine levels. (**A**) Experimental setup and hierarchical clustering of inflammatory mediator expression in all tested SF samples. (**B**-**D**) Markers with significant elevation in sJIA SF compared to JIA SF (**B**), non-significant elevation in just JIA SF (**C**) and only by-trend differential expression in JIA compared to sJIA SF (**D**). Orange bars in each plot indicate healthy donor serum levels (10-90^th^ percentile) as a reference. Data were analyzed by Mann-Whitney U test. (**E**) Spearman r values for association of indicated cytokines with Th cellular ICOS expression fold change as induced upon cell cultures in 3% JIA or sJIA SF. Colored bars indicate significant association. (**F**, **G**) Exemplary correlation plots of S100A9 and GM-CSF SF levels with Th cellular ICOS expression fold change induced upon cell cultures in 3% (F) or 6% (G) JIA or sJIA SF. sJIA and JIA SF donors were indicated by circles and squares respectively. (**H**, **J**) Spearman r values for the association of indicated cytokines with Tph (**H**) and Tfh (**J**) IFNγ expression fold change as induced upon cell cultures in 3% JIA or sJIA SF. Colored bars indicate significant association. (**I**, **K**) Exemplary correlation plots of GM-CSF SF levels with Tph (**I**) and Tfh (**K**) IFNγ expression fold change induced upon cell cultures in 3% or 6% JIA or sJIA SF. Circles indicate individual sJIA and squares individual JIA SF donors. * = p<0.05, ** = p<0.01; *** = p<0.001

Next, we tested associations of SF inflammatory mediator expression with several of the SF-induced Th cell phenotypes, which revealed significant differences depending on whether cells were cultured in a JIA or sJIA SF environment (**Fig. 3E-K**). We observed Th cellular ICOS expression, which was significantly increased upon cell culture in sJIA SF but not JIA SF conditioned medium (**Fig. 1D**, **Fig. 2C**) to strongly associate with several markers that were significantly overexpressed in sJIA compared to JIA SF (S100A9, E-selectin, IL-18, GM-CSF), but also to link with inflammatory mediator levels in SF with no significant difference between JIA and sJIA (CCL11, MMP-7, IL-1Ra; **Fig. 3E-G**). Beyond this, we found IFNγ expression from cells with a Tph-phenotype induced upon culture in JIA or sJIA SF (**Fig. 2H**) to best associate with SF levels of IL-1Ra, IL-8, GM-CSF, MMP-7, IL-1β and S100A9 (**Fig. 3H, I**), of which GM-CSF, IL-1β and S100A9 were significantly overexpressed in sJIA compared to JIA SF (**Fig. 3B**). Similarly, IFNγ expression from cells with a Tfh-phenotype induced upon culture in JIA or sJIA SF (**Fig. S2D**) was significantly associated with IL-1Ra, GM-CSF, S100A9 and IL-1β SF levels (**Fig. 3J, K**). Among these, GM-CSF, IL-1β and S100A9 were significantly overexpressed in sJIA compared to JIA SF (**Fig. 3B**). Furthermore, we observed increases in CXCR3^neg^CCR6^pos^ Th cell frequencies upon culture in JIA SF and expansion of CXCR3^pos^CCR6^pos^ Th cells in sJIA SF conditioned medium to associate with multiple parameters (**Fig. S3**).

### Spiking JIA SF with selected inflammatory cytokines can revert some T cellular phenotypes towards those observed in sJIA SF cultures

Association studies as described above (**Fig. 3E-K**) can help to pinpoint factors potentially contributing to the observed cell phenotypes. However, such analysis can at best suggest candidate markers and do not allow to argue for a true causal link. Therefore, we set out to better understand, whether some of the identified associations may indeed reflect a direct impact of inflammatory markers on specific cell phenotypes. To investigate this, we spiked selected recombinant cytokines (IL-1β, IL-18, GM-CSF) based on two criteria: 1) these cytokines were significantly increased in sJIA compared to JIA SF (**Fig. 3B**) and 2) they showed significant associations with the reported cell phenotypes into JIA SF samples (**Fig. 3E-K**. Recombinant cytokine concentrations were chosen according to median levels quantified in sJIA SF and cytokines were added either alone or in combination (**Fig. 4A**).

**Figure 4.**
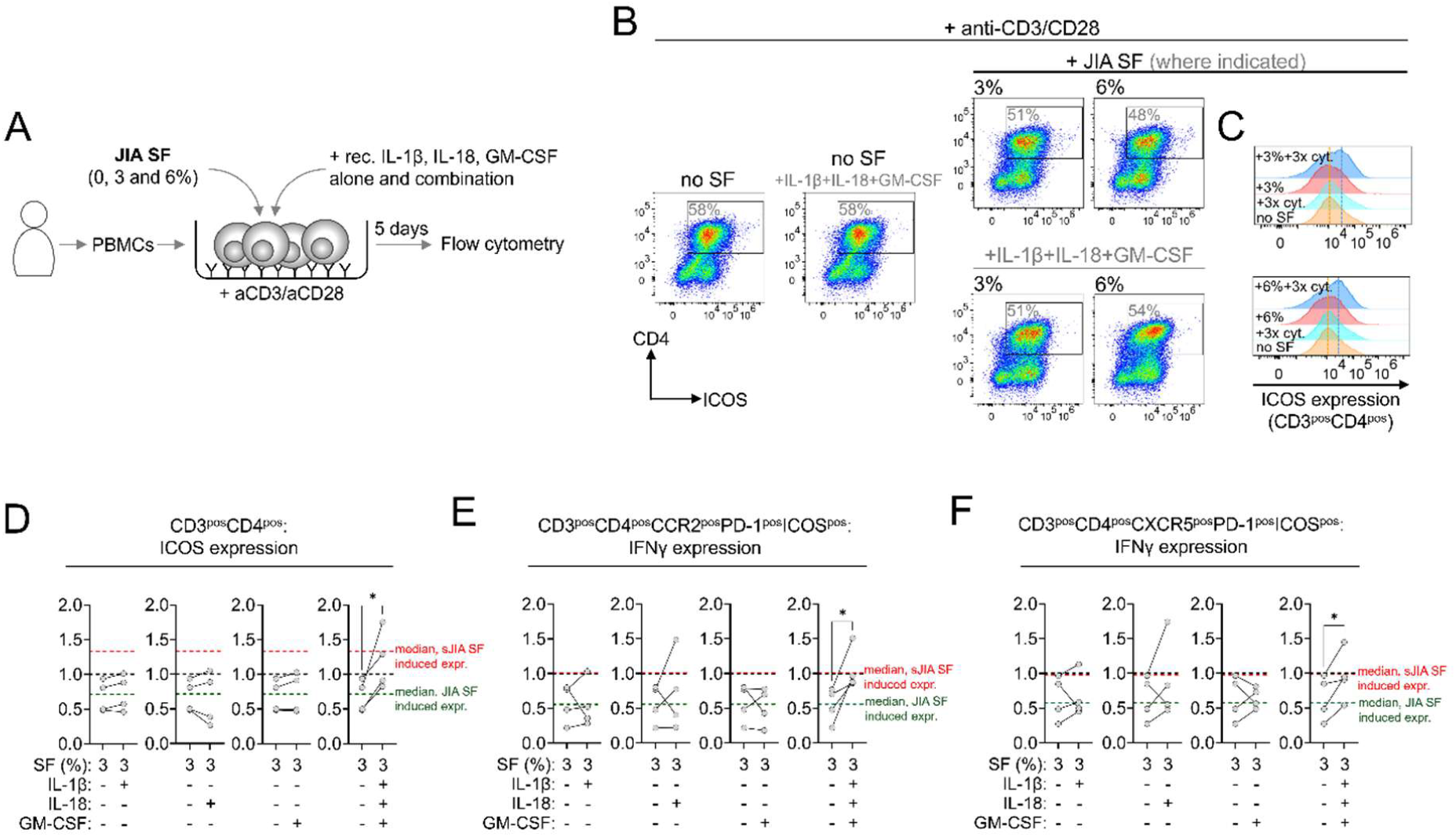
Spiking JIA SF with selected inflammatory cytokines can revert some T cellular phenotypes. (**A**) Experimental setup. (**B**, **C**) Exemplary pseudocolor flow cytometry and histogram plots (**C**) of CD4 T cellular ICOS expression following five days of HD PBMC culture with anti-CD3/CD28 stimulation and JIA SF (both 3 and 6%) conditioned medium and addition of indicated recombinant cytokines. (**C**) Dashed vertical lines in histogram plots indicate peak Th cellular ICOS expression with just anti-CD3/CD28 stimulation (orange) and upon culture with JIA SF (3%: upper panel, 6%:lower panel) and addition of recombinant IL-1β, IL-18 and GM-CSF (blue). (**D**-**F**) Cumulative data (n=4 independent JIA SF donors) of Th cellular ICOS (**D**), Tph (**E**) and Tfh (**F**) IFNγ expression fold change induced by the indicated cell culture conditions. Dashed lines in all plots indicate expression levels induced by just anti-CD3/CD28 stimulation (1.0, black) and median fold change induced upon culture in 3% JIA (green) or sJIA SF (red). Data were analyzed by paired t-test. * = p<0.05

When assessing Th cellular ICOS expression in these experiments, which was significantly decreased upon cell culture in JIA SF compared to just anti-CD3/CD28 stimulation or culture in sJIA conditioned medium in previous analysis (**Fig. 1D**, **Fig. 2C**), simultaneous addition of recombinant IL-1β, IL-18 and GM-CSF to JIA SF cultures significantly increased Th cellular ICOS expression (**Fig. 4B, C**). This finding suggests that the addition of these cytokines partially mimics the expression phenotype observed when cells were cultured in sJIA SF conditioned medium (**Fig. 4C**; **Fig. 1D**; **Fig. 2C**). We did not observe this increase upon addition of each recombinant cytokine to JIA SF individually (**Fig. 4B, C**). Moreover, we observed that a combination of recombinant IL-1β, IL-18 and GM-CSF also modified IFNγ expression from Tph (**Fig. 4E**) and Tfh cells (**Fig. 4F**) in cell cultures with JIA SF towards levels as induced by sJIA SF conditioned medium in our previous experiments (**Fig. 2H** and **Fig. S2D**). In contrast, recombinant cytokine addition to JIA SF did not result in any significant frequency changes of CXCR3^neg^CCR6^pos^ and CXCR3^pos^CCR6^pos^ Th cell subsets (**Fig. S5**), which we previously observed to be differentially expanded in cell cultures with either JIA or sJIA SF conditioned medium (**Fig 1G, I**).

## Discussion

Joint inflammation in context of JIA is frequently treated by intra-articular injection of steroids, which can give access to synovial fluid and cells. In context of sJIA, this is done only in rare occasions and limited to cases, where systemic immunosuppressive therapy repeatedly fails to control synovial inflammation.

Regardless of the underlying JIA disease category, the synovial environment in context of ongoing articular inflammation is highly inflammatory with multiple proteins, cyto-and chemokines abundantly expressed, in part in levels exceeding those in serum or plasma by far(26, 27). While inflammatory mediator profiles in SF are overlapping between systemic, oligo-and polyarticular JIA(26, 27), some differential features(26, 27) and distinct cell populations(13) as well as differing responsiveness to inflammatory challenge have been reported(28).

Benefitting from the rare opportunity of having access to sJIA synovial biomaterial, in the present study we report a series of functional experiments, which demonstrate differential Th cell polarization and activation, including Tf/ph cell phenotypes upon exposure to either systemic or poly-/oligo (“polygo”) JIA SF. These data, linking to sites of ongoing tissue-borne inflammation, complement our previous findings from the sJIA periphery, which demonstrated a wiring of sJIA naïve Th cell differentiation towards a Tf/ph (precursor) phenotype(17). Importantly, in their interpretation - particularly with respect to Tph prevalence in sJIA and role of IL-1β/IL-18 signaling - our present data also complement a deep-phenotyping and single cell RNA-sequencing study on sJIA *versus* JIA synovial Th cells that was conducted in parallel (22).

When testing the hypothesis of differential inflammatory T cell activation upon exposure to either a JIA or sJIA SF inflammatory matrix, one set of data from our study suggests a striking and differential expansion of CXCR3^neg^CCR6^pos^ Th cells upon healthy donor T cell stimulation (anti-CD3/CD28) in context of JIA SF, compared to an expansion of CXCR3^pos^CCR6^pos^ Th cells in cell cultures involving sJIA SF, instead. CCR6 is a surface marker, that has been reported to be expressed on both Th17 as well as non-classical Th1 (Th17.1) cells, yet the latter also expressing CXCR3(23). This may imply JIA SF to preferentially promote expansion of Th17 cells, whereas sJIA SF rather drives the expansion of a non-classical Th1 (Th17.1) cell population, which appears significantly depleted upon cell cultures in JIA SF. CXCR3^neg^CCR6^pos^ Th17 cells are reported as prominent producers of IL-17(23), which we also observed from our data when staining for intracellular IL-17A expression in this subset. In contrast, Th17.1 cells are regarded as strong producers of IFNγ, but not IL-17(23). In this respect, the observed expansion of Th17.1 cells upon T cell stimulation in the context of sJIA SF somewhat contrasts our data on cytokine expression into cell culture supernatants, as here we find IL-17A release to be markedly promoted over IFNγ expression by sJIA SF in all tested cell culture conditions. In contrast, JIA SF promotes some IL-17A release, while depleting IFNγ expression, which may indeed reflect a JIA SF driven Th17 expansion. Importantly, in our experiments both the expansion of CXCR3^neg^CCR6^pos^ as well as IL-17A expressing CXCR3^neg^CCR6^pos^ Th cells are significantly reduced particularly upon IL-1R inhibition, which fits a prominent role of both IL-1α and IL-1β signaling in Th17 differentiation and expansion(29).

In the context of these data, several other studies already reported elevated numbers of IL-17 producing Th cells to be present in cell preparations obtained from JIA SF, which have been considered either bona fide Th17 or Th17.1 cells (30, 31). Contrasting these studies as well as our data linked to polygo JIA SF, a study by Julé and colleagues described an overly prominent Th1 (CXCR3^pos^IFNγ^pos^) and only moderate Th17 polarization of the memory T cell compartment within oligo JIA SF(32). Yet, these data cannot be directly compared with the results from our experimental approach, as Julé et al assessed cell phenotypes within SF(32), while we report on the impact of the JIA SF inflammatory environment on healthy donor Th cell differentiation and activation.

As main hypothesis of the present study, however, we aimed to assess the impact of an sJIA versus JIA inflammatory environment on Tf/ph cells, as from our previous data it remained unclear whether the reported differences in T cell differentiation in sJIA may occur due to sJIA-specific genetic programs within T cells, or are a secondary effect induced by the disease specific inflammatory environment. Obtaining biomaterial from both JIA as well as sJIA inflamed joints as an inflammatory hot-spot allowed to test for an impact of a matrix enriched for inflammatory mediators on T cell biology.

Collectively, our flow-cytometry data suggest T cell stimulation (anti-CD3/CD28) in the presence of neither JIA nor sJIA SF to significantly promote expansion of Th cells with a Tfh-(ICOS^pos^PD-1^pos^CXCR5^pos^) or Tph-phenotype (ICOS^pos^PD-1^pos^CCR2^pos^). However, cell cultures in sJIA but not JIA SF conditioned medium significantly sustained Tph-phenotypes when those were induced by additional cell stimulation with recombinant cytokines. This effect was even more pronounced when analyzing IFNγ-expression from Tf/ph cells. This appeared significantly reduced upon T cell stimulations in the presence of JIA SF, but remained unaffected upon cell culture in sJIA SF conditioned medium.

Beyond, cytokine release and gene expression data from our experiments recapitulate differential IFNγ expression upon cell cultures in JIA *versus* sJIA SF, but also imply more robust T cellular IL-21 and c-MAF expression upon cell cultures in the presence of sJIA SF. Conversely, T cell stimulation in the presence of JIA SF with or without the addition of recombinant cytokines resulted in a marked increase in CXCL-13, Bcl-6 and Blimp-1 (*PRDM1*) expression. While both IL-21 and CXCL-13 are considered as defining cytokines of both Tfh as well as Tph cells, c-MAF and particularly Blimp-1 are transcription factors driving Tph development. In contrast, Blimp-1 is typically downregulated in Tfh cells, which is hallmarked by increased expression of the transcription factor Bcl-6, instead(18).

Indeed, increased Th cellular ICOS expression, as observed throughout in T cell stimulations in the presence of sJIA but not JIA SF in our flow cytometry analysis, may directly link with elevated/sustained effector cytokine expression as observed for IFNγ and IL-21(33). Yet, further interpreting flow cytometry, cytokine and gene expression data altogether remains complex. A perseverance of Th cells with a Tph-phenotype with prominent IFNγ, but also IL-21 and c-MAF expression, can argue for sJIA SF to selectively promote a self-sustaining (IL-21, c-MAF) Tph subset with the potential to provide strong B cell support towards plasmablast generation (IFNγ)(25). In contrast, T cell stimulation in the presence of JIA SF may support development of a Tf/ph intermediate phenotype (considering simultaneous increase in both Bcl-6 and Blimp-1 expression) with strong B cell recruiting capabilities (CXCL-13)(25).

From our experiments, we have some indications on the specific inflammatory matrix impact on T cell activation in either JIA or sJIA SF context. A differential impact of the JIA *versus* sJIA SF matrix on Th cellular ICOS expression is somewhat supported by blocking prominent inflammatory axis (IL-6 and IL-1(β) signaling) in respective stimulation experiments. IL-6 and IL-1α/β are abundantly present in both JIA and sJIA SF, with IL-1β levels significantly elevated in sJIA compared to JIA SF. Blocking either the IL-1 or IL-6 receptor, or neutralizing IL-1β affects the JIA or sJIA SF induced ICOS expression phenotypes by trend. In these lines, particularly IL-6R *cis-*signaling has already been reported to be absolutely required for Th cellular ICOS expression (34), while IL-1 signaling to human Th cells benefits multiple effector functions and cytokine expression across different Th cell lineages, but (at least for stimulation times of 36h or less) has not been reported to impact ICOS expression(35). Furthermore, our association studies and experiments involving recombinant cytokine spiking hint at a beneficial and combined impact of elevated IL-1β, IL-18 and GM-CSF levels in sJIA SF on supporting Tph-phenotypic features as observed with T cell stimulations in sJIA SF conditioned medium, such as increased Th cellular ICOS as well as IFNγ expression from ICOS^pos^PD-1^pos^CCR2^pos^ and ICOS^pos^PD-1^pos^CCR2^pos^ Th cell subsets. Of note, all three cytokines have already been associated with beneficial effects on Tf/ph polarization(36, 37), activation(37, 38), expansion (IL-18;(39)) and general Th eff function (IL-1; (35)). In contrast, in these experiments we observed no impact of recombinant cytokine spiking on expansion of a Th17.1 (CXCR3^pos^CCR6^pos^) subset as observed upon T cell stimulations in sJIA SF conditioned medium.

Importantly, our study data require to be interpreted in light of several limitations. One obvious draw-back is the limited number of sJIA SF samples, that have been included. This is due to the rare availability of this particular material from this group of patients at our end. Next, SF represents a highly variable matrix, also considering different medications both JIA and sJIA patients received (in part) at time of sampling. In order to buffer for some of this variability our cell stimulation experiments include PBMCs and T cells of just a single healthy donor as a constant throughout all experiments, which allowed us to better study SF impact on Th cell biology. T cell targeted stimulation was always performed in context of whole PBMCs, in order for the quantified effects on Th cell level to reflect SF-impact on complex cellular networks, and not pre-sorted cells only. Beyond, our experiments are very much hypothesis driven and were thus restricted to analysis of only specific cell surface, cytokine or transcription factor expression; we are aware, that this excludes several other possible traits of SF-impact on Th cell biology. Yet, we also monitored SF-impact on CD8^pos^ cytotoxic T cells, which, however, is beyond the scope of the present study. Finally, our study also excludes potential kinetic trajectories of T cell activation and development, as all analysis for surface marker, cytokine or gene expression were performed after five days of culture.

Despite these limitations, we believe our study to illustrate, that JIA and sJIA SF can drive distinct Th cell subset polarization or activation, including sustaining different Tf/ph cell phenotypes and effector functions. These findings complement our earlier observations from sJIA peripheral blood and demonstrate the impact of the SF inflammatory matrix on immune cell activation.

### Contributors

JD, HM and CK designed the study. SS, AS and SF performed experiments. SS, AS, SF and CK collected experimental data. CB, IC, TH, SH, GP, GT, MP, CP, CH, HW, DW and DF collected patient samples and data. SS, JD, SF, HM and CK analyzed data. SS, AS and CK drafted the manuscript, and all authors revised and approved the final draft.

**COI statement:** CB received consultancy fees from Sobi and Novartis and speaker fees from GSK. MP received consultancy fees from Sobi and Novartis. CH has received honoraria (lecture fees) from Novartis; HW has received honoraria (lecture fees) from Novartis and Takeda, and travel support from Octapharma and CSL-Behring; DF received speaker fees/honoraria from Chugai-Roche, Novartis and SOBI as well as research support from Novartis, Pfizer and SOBI. HM received honoraria (lectures fees) and travel support from Novartis. No other disclosures relevant to this article were reported. CK has received consulting fees from Novartis and Swedish Orphan Biovitrum (SOBI) (< $10,000 each) and received research support from Novartis (> $10,000).

**Sources of support:** AS received support from the Muenster University medical faculty MedK program. JD was supported by the Interdisciplinary Center for Clinical Research (IZKF) Würzburg (Bridging Program, Z-3/BC-13). GP was supported by the Italian Ministry of Health with “Current Research funds”. HM received funding from the Federal Ministry of Education and Research (BMBF; Advanced Clinician Scientist-Program INTERACT; 01EO2108) embedded in the Interdisciplinary Center for Clinical Research (IZKF) of the University Hospital Würzburg. HM also receives support from the German Research Foundation (MO 2160/4-1, HM). CK receives support from the German Research Foundation (DFG, grant number KE 2026 1/3).

## Ethics approval

The study was reviewed by the Research Ethic Committee of the University of Münster (2015-670-fS, covering sample collection in Münster and Sendenhorst) as well as the Ethical Committee of Ospedale Pediatrico Bambino Gesù in Rome (2333/OPBG 2020) and conducted in accordance with the Declaration of Helsinki. All patients or care givers signed written informed consent.

## Data availability statement

Study data are available from the corresponding author upon reasonable request.

