## supplementary methods and data for "The systemic JIA synovial fluid environment supports development and prevalence of specific inflammatory T helper cell phenotypes"

*Study patients and samples*

Systemic JIA patients included in this study were followed at the Division of Rheumatology, Ospedale Pediatrico Bambino Gesù, IRCCS, Rome, Italy. JIA patients' samples were obtained from the Clinic for Pediatric and Adolescent Rheumatology, Northwest German Center for Rheumatology, St. Josef Stift Sendenhorst, or the Department of Pediatric Rheumatology & Immunology, University Hospital Münster, Münster, Germany. The study was reviewed by the Research Ethic Committee of the University of Münster (2015-670-fS, covering sample collection in Münster and Sendenhorst) as well as the Ethical Committee of Ospedale Pediatrico Bambino Gesù in Rome (2333/OPBG 2020) and conducted in accordance with the Declaration of Helsinki. All patients or care givers signed written informed consent. During routine joint aspiration, cell-free synovial fluid was obtained from all study patients and stored at -20°C until use. For all cell stimulation experiments, freshly prepared PBMCs prepared from blood of a single healthy adult donor (several donation time points) were used.

*Human peripheral blood mononuclear cell (PBMC) isolation and culture*

PBMCs were isolated by density-gradient centrifugation (Pancoll, density: 1.077 g/ml, PAN-Biotech GmbH, Germany) from peripheral blood samples collected from a single healthy donor. For PBMC activation/differentiation experiments PBMCs freshly isolated from a single healthy donor were directly used in experiments. PBMCs were diluted to $5 \times 10^5$ cells x $ml^{-1}$ in supplemented RPMI 1640 medium and plated at a density of $5 \times 10^4$ cells / well in 96-well U-bottom suspension culture plates (Greiner Bio-One, Frickenhausen, Germany) coated with α-human CD3 (5 µg x $ml^{-1}$, clone: OKT3, BioLegend, San Diego, CA, USA). Cells were cultured for 5 days (37°C, 5% $CO_2$) with or without cell-free JIA or sJIA synovial fluid at final concentrations of 3% or 6% and in the presence of combinations of the following cytokines, stimuli, or blocking agents, as indicated and detailed in the respective results: α-human CD28 (2.5 µg x $ml^{-1}$, clone: CD28.2, BioLegend), recombinant human IL-1β (150 ng x $ml^{-1}$, BioLegend), recombinant human IL-2 (100 U x $ml^{-1}$, PeproTech), recombinant human IL-6 (5 ng x $ml^{-1}$, PeproTech), recombinant human IL-12 (5 ng x $ml^{-1}$, PeproTech), recombinant human IL-18 (10 ng x $ml^{-1}$, Invivogen, San Diego, CA, USA), recombinant human IL-21 (20 ng x $ml^{-1}$, BioLegend), recombinant human GM-CSF (50 pg x $ml^{-1}$, PeproTech), anti-human-IL-6R mAb (tocilizumab, 1000nM), recombinant human IL-1Ra (anakinra, 200 ng x. $ml^{-1}$) and anti-human-IL-1β (canakinumab, 1000 ng x. $ml^{-1}$).

*Flow cytometry*

Before flow cytometry analysis, PBMCs or previously naïve Th cells were stimulated by incubation with phorbol 12-myristate 13-acetate (PMA) and ionomycin (eBioscience™ Invitrogen™ Cell Stimulation Cocktail, Thermo Fisher Scientific Inc.) in the presence of Brefeldin A and Monensin (eBioscience™ Invitrogen™ Protein Transport Inhibitor Cocktail, Thermo Fisher Scientific Inc.) for 5 hours at 37 °C, or left unstimulated. For cell surface marker staining, $1 \times 10^5$ cells were incubated with the respective antibody at a concentration of 1 µl / 100 µl sample in a buffer containing 1 x phosphate-buffered saline (PBS; Merck KGaA, Darmstadt, Germany), 0.1 % bovine serum albumin (BSA; Carl Roth GmbH, Karlsruhe,

Germany), 2 mM ethylenediaminetetraacetic acid (EDTA; Carl Roth GmbH), and 0.05 % sodium azide (Merck KGaA) for 30 minutes at 4 °C in the dark.
The following antibodies were used for T cell surface marker staining: α-human CD3-APC-Cy7 (clone: OKT3, BioLegend), α-human CD4-APC (clone: OKT4, BioLegend), α-human CD4-BV510 (clone: OKT4, BioLegend), α-human CD183/CXCR3-APC-Cy7 (clone: G025H7, BioLegend), α-human CD185/CXCR5-PE-Dazzle (clone: J252D4, BioLegend), α-human CD196/CCR6-BV650 (clone: G034E3, BioLegend), α-human CD278/ICOS-FITC (clone: C398.4A, BioLegend), and α-human CD279/PD-1-BV650 (clone: EH12.2H7, BioLegend).
For the subsequent staining of intracellular cytokines and transcription factors, the cells were fixed and permeabilized by incubation in FixPerm Buffer (eBioscience™ Fixation/Permeabilization Concentrate, eBiosciene™ Fixation/Permeabilization Diluent, Thermo Fisher Scientific Inc., USA) for 30 minutes at room temperature. For intracellular staining, $1x10^5$ cells were incubated with the respective antibody at a concentration of 1 µl / 100 µl sample in 1 x Permeabilization Buffer (eBioscience™, Thermo Fisher Scientific Inc., USA) for 30 minutes at 4 °C in the dark. The following antibodies were used for intracellular staining: α-human IFNγ-BV421 (clone: 4S.B3, BioLegend, USA), α-human IL-17A-PE (clone: BL168, BioLegend, USA), α-human IL-21-PE (clone: 3A3-N2, BioLegend, USA). Fluorescence intensities were measured with CytoFLEX S (Beckman Coulter, USA). For compensation and gating, fluorescent beads and unstained samples were used, respectively. Measurement settings were not changed throughout the study. Cytometric data were analyzed by FlowJo (version: v10.0.8, Becton, Dickinson and Company, USA). Expression levels of cytokines and transcription factors were assessed as geometric mean fluorescence intensities (MFIs). Changes in expression were calculated as fold change compared to respective MFIs of cells stimulated with antiCD3/CD28 alone. Changes in population frequencies were calculated based on fold change in percentage of respective absolute cell numbers in total single cells, compared to stimulated with antiCD3/CD28 alone.

*Bead array assay (Luminex)*

Reagents for multiplexed quantification of indicated markers in JIA and sJIA synovial fluids or culture supernatants were purchased from R&D Systems (Minneapolis, OH, USA). Reagents and samples were prepared according to the manufacturer's instructions (R&D Systems). All samples were measured in serial dilution (1:2, 1:4, 1:8, 1:16, 1:32, 1:64). Data acquisition and analysis were performed on a MAGPIX instrument (Merck Millipore, Darmstadt, Germany) using xPONENT v4.2 software (Luminex).

*Gene expression*

Cells were harvested and resuspended in DNA/RNA Shield (Zymo Research, Freiburg, Germany), and RNA was isolated using the Quick-RNA™ Whole Blood Kit according to the manufacturer's protocol. RNA was transcribed using RevertAid H Minus Reverse Transcriptase (Thermo Fisher, Karlsruhe, Germany) and used as template for qRT-PCR. qRT-PCRs were prepared using KAPA SYBR FAST qPCR Kit (Merck, Darmstadt, Germany) according to the manufacturer's instructions and cycling was performed on a CFX 384 Real-Time System (Bio-Rad, Feldkirchen, Germany) by the Core Facility Genomics at Muenster University medical faculty. Respective oligonucleotides are listed in **table S1**.

*Data analysis*

Statistical analyses was performed using GraphPad Prism software (version 10.2.2, GraphPad Software Inc., USA). The statistical tests applied for individual data sets are indicated in the respective figure legends. $P<0.05$ was considered as statistically significant. Synovial fluid inflammatory marker data were analyzed for unsupervised clustering using correlation distance and ward.D2 linkage by the pheatmap R-package and Rstudio (RStudio Team (2015). RStudio: Integrated Development for R. RStudio, Inc., Boston, MA http://www.rstudio.com/).

**Table S1.** Oligonucleotides for qRT-PCR

| **Gene** | **forward** (5'-3') | **reverse** (5'-3') |
|---|---|---|
| *BCL6* | atgccagtgatgttcttctcaa | aaggttgcatttcaactggtct |
| *PRDM1* | gactttgcagaaaggcttcact | gaatcacatgaagggcatgtt |
| *IL21* | aggaaaccaccttccacaaa | gaatcacatgaagggcatgtt |
| *IFNG* | gcatcgttttgggttctcttg | agttccattatccgctacatctg |
| *CXCL13* | tctctgcttctcatgctgct | tcaagcttgtgtaatagacctcca |
| *MAF* | ccgtcctctcccgagttttt | tgctggggcttccaaaatgt |

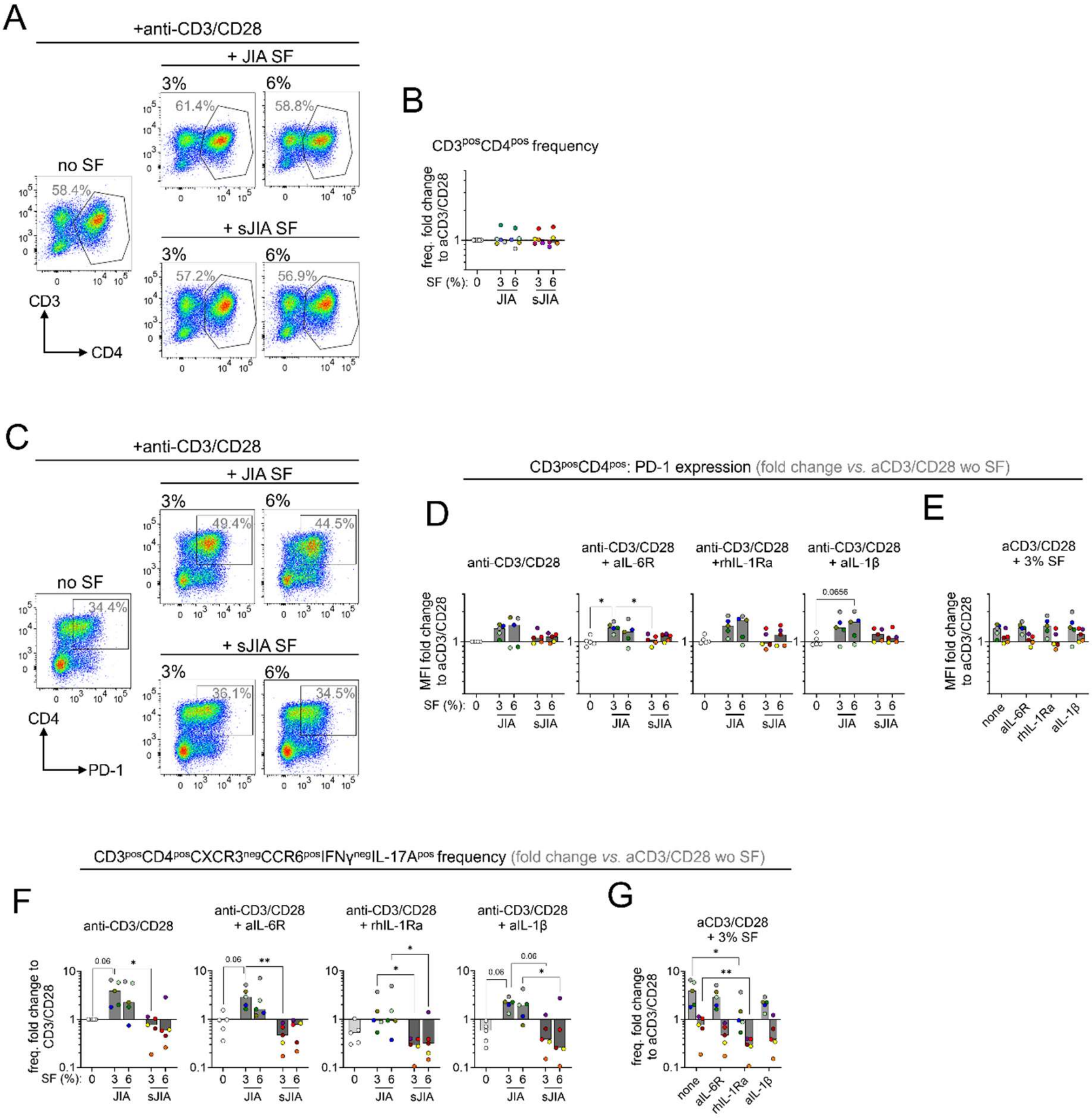


**Figure S1.** Differential Th cell activation and subset expansion upon HD PBMC culture in JIA or sJIA SF conditioned medium. (**A**, **B**) Exemplary pseudocolor flow cytometry plots and cumulative data (**B**; n=5 independent JIA SF donors; n=5 sJIA independent SF donors) of CD3$^{pos}$CD4$^{pos}$ T cell frequency following five days of HD PBMC culture with anti-CD3/CD28 stimulation and JIA or sJIA SF (both 3 and 6%) conditioned medium. Changes in population frequencies were calculated based on fold change of absolute cell numbers in total single cells, compared to cells stimulated with anti-CD3/CD28 alone. (**C**) Exemplary pseudocolor flow cytometry plots of CD4$^{pos}$PD-1$^{pos}$ cells following five days of HD PBMC culture with anti-CD3/CD28 stimulation and JIA or sJIA SF (both 3 and 6%) conditioned medium. (**D**, **E**) Cumulative data (n=5 independent JIA SF donors; n=5 sJIA independent SF donors) of CD4 T cellular PD-1 expression fold change induced by the indicated cell culture conditions, with or without additional *in vitro* drug treatment (anti-IL-6R, tocilizumab; recombinant human IL-1Ra, anakinra; anti-IL-1b, canakinumab). Changes in expression were calculated as fold change compared to mean fluorescence intensities of cells stimulated with only anti-CD3/CD28. (**F**, **G**) Cumulative data (n=5 independent JIA SF donors; n=5 sJIA independent SF donors) of IL-17A expressing CCR6$^{pos}$ CD4 T cell frequency fold change induced by the indicated cell culture conditions, with or without additional indicated *in vitro* drug. Changes in frequency were calculated as fold change compared to cell expansion stimulated with only anti-CD3/CD28. All data were analyzed by Friedman test for multiple paired non-parametric observations, followed by Dunn's multiple comparison test. * = p<0.05, ** = p<0.01

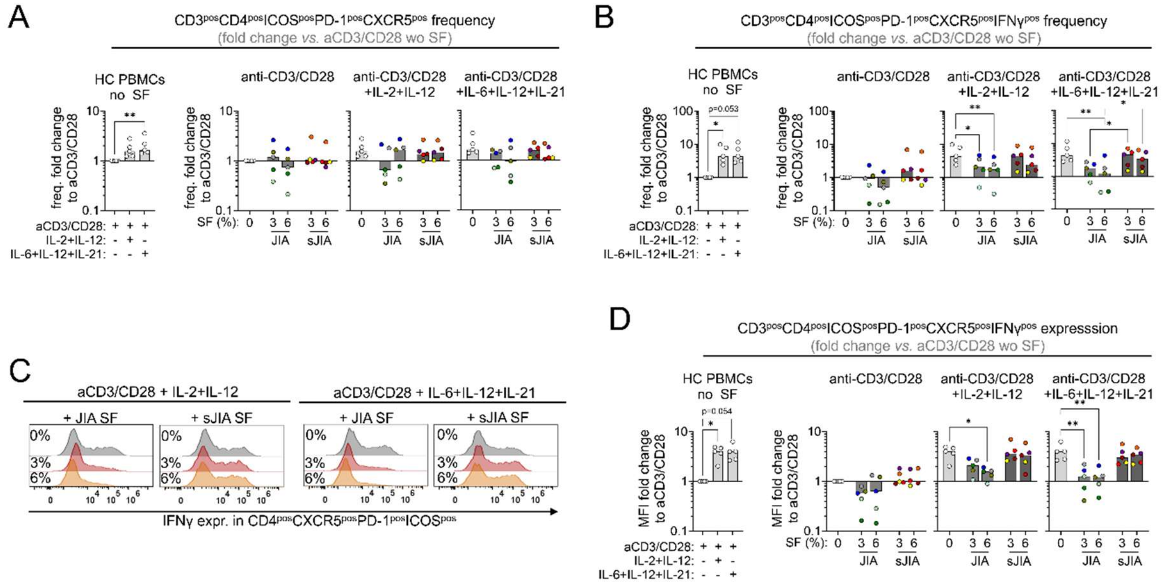


**Figure S2.** Systemic JIA but not JIA SF preserves a recombinant cytokine induced T follicular helper cell (Tfh) phenotype. (**A**, **B**) Cumulative data (n=5 independent JIA SF donors; n=5 sJIA independent SF donors) of Tfh (**A**) and IFNγ expressing Tfh expansion (**B**) fold change induced by the indicated cell culture conditions. Changes in population frequencies were calculated based on fold change of absolute cell numbers in total single cells, compared to cells stimulated with anti-CD3/CD28 alone. (**C**, **D**) Exemplary histogram plots (**C**) and cumulative data (n=5 independent JIA SF donors; n=5 sJIA independent SF donors) of Tfh IFNγ expression fold change (**D**) induced by the indicated cell culture conditions. (**C**, **H**) Changes in expression were calculated as fold change compared to mean fluorescence intensities of cells stimulated with only anti-CD3/CD28. All data were analyzed by Friedman test for multiple paired non-parametric observations, followed by Dunn's multiple comparison test. * = p<0.05, ** = p<0.01

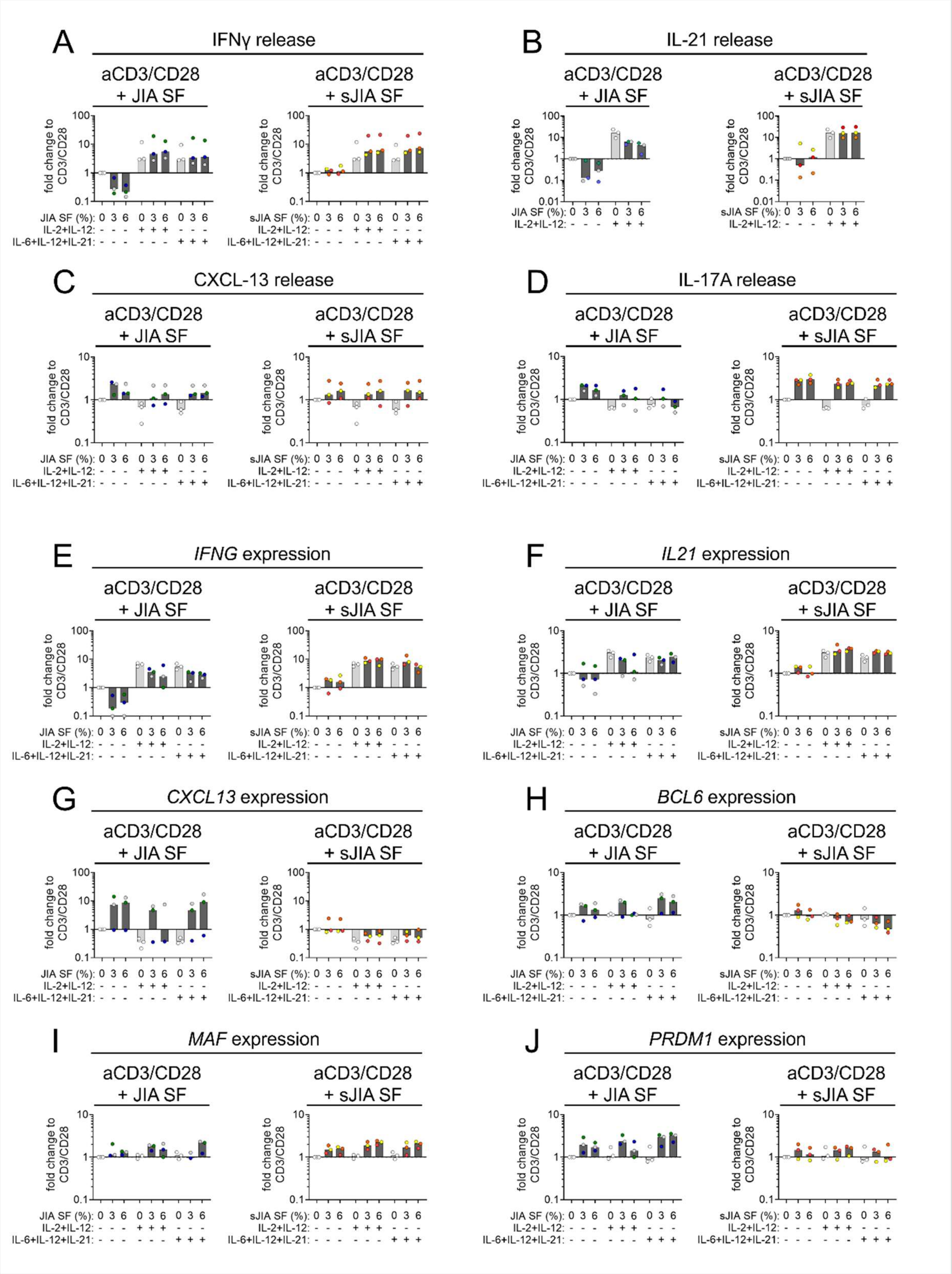


**Figure S3.** Cytokine release and gene expression analysis of T cell stimulations in cell cultures with JIA or sJIA SF. (**A-D**) Cytokine release into cell culture supernatants was assessed by multiplexed bead array assay. Only netto expression over respective cytokine backgrounds already present in SF was considered for data analysis. IL-21 release was not quantified in cell cultures spiked with recombinant IL-21 (**B**). (**E-J**) Expression of selected genes in cell cultures receiving T cell stimulation (anti-CD3/CD28, with or without the addition of recombinant cytokines) in the presence of either JIA or sJIA SF was assessed by qRT-PCR. (**A**-**J**) Changes in expression were calculated as fold change compared to gene or protein expression by cells stimulated with anti-CD3/CD28 alone.

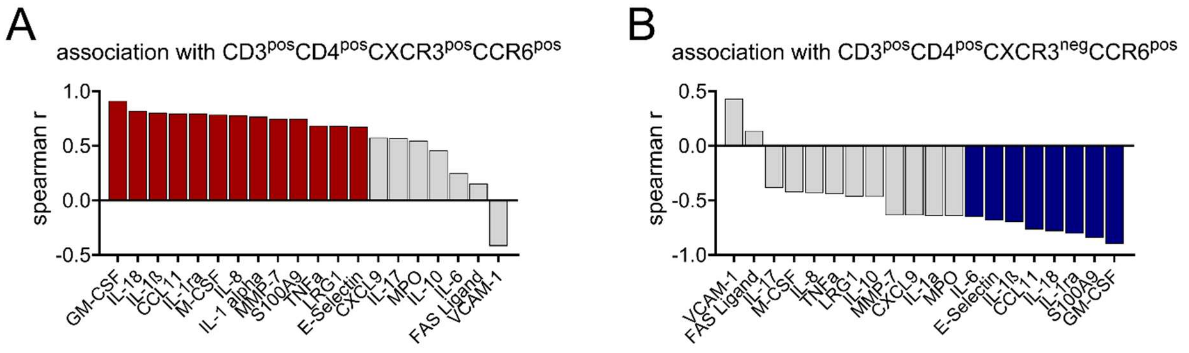


**Figure S4.** Association of inflammatory mediator levels in SF and inflammatory Th cell phenotypes. (**A**, **B**) Spearman r values for association of indicated cytokines with CD3$^{pos}$CD4$^{pos}$CXCR3$^{pos}$CCR6$^{pos}$ (**A**) and CD3$^{pos}$CD4$^{pos}$CXCR3$^{neg}$CCR6$^{pos}$ (**B**) T cell expansion fold change as induced upon cell cultures in 3% JIA or sJIA SF. Colored bars indicate significant association.

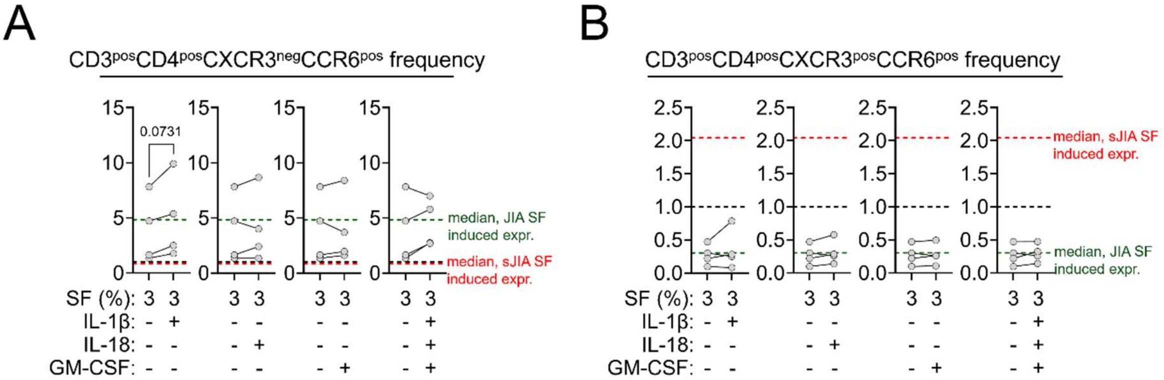


**Figure S5.** Impact of inflammatory cytokine spiking into JIA SF on specific inflammatory T cell phenotypes. (**A**, **B**) Cumulative data (n=4 independent JIA SF donors) of CD3$^{pos}$CD4$^{pos}$CXCR3$^{pos}$CCR6$^{pos}$ (**A**) and CD3$^{pos}$CD4$^{pos}$CXCR3$^{neg}$CCR6$^{pos}$ (**B**) T cell expansion fold change induced by the indicated cell culture conditions. Dashed lines in all plots indicate expression levels induced by just anti-CD3/CD28 stimulation (1.0, black) and median fold change induced upon culture in 3% JIA (green) or sJIA SF (red). Data were analyzed by paired t-test. * = p<0.05
